# Developmental Olfactory Experience Dissociates Hedonic Valence from Exploratory Arousal in *Drosophila melanogaster*

**DOI:** 10.64898/2026.04.29.721699

**Authors:** Matías Alemán, Lautaro Alejandro Duarte, Fernando Federico Locatelli, Nicolás Pírez

## Abstract

Insects rely on olfactory cues to navigate complex environments, with many innate behaviors governed by evolutionary hardwired neural circuits. However, the extent to which early-life sensory experience can recalibrate these innate responses remains a subject of intense debate. Here, we investigate how chronic exposure to odorants of varying innate valences during development and early adulthood modulates olfactory preference and exploratory drive in *Drosophila melanogaster*. Using a high-resolution two-choice assay, we demonstrate a fundamental functional dissociation in olfactory plasticity: while the innate hedonic valence of most odorants remains remarkably resilient to developmental environmental manipulation, early-life experience profoundly reprograms exploratory dynamics. Specifically, chronic exposure to propionic acid and benzaldehyde induced sex-specific shifts in locomotor arousal and trap-entry decisions without altering the intrinsic hedonic valuation of the stimuli. Conversely, general exploratory drive toward 1-octanol and isoamyl acetate remained rigidly hardwired, although 1-octanol exhibited subtle, experience-dependent habituation in odor preference. This resilience of innate valence suggests that the olfactory circuit actively prioritizes functional stability to ensure that critical ecological cues remain reliably encoded. Our findings reveal that *Drosophila* employs a modular adaptive strategy to integrate chronic sensory information: unreinforced early-life experience selectively reconfigures motor reactivity to scale navigational intensity to familiar landscapes, while leaving primary sensory-driven valences largely intact.

## Introduction

Olfaction is a fundamental sensory modality for insects, providing essential information about food sources, predators, mating opportunities, and oviposition sites. In the fruit fly *Drosophila melanogaster*, innate olfactory responses are mediated by well-characterized neural circuits, making this species an ideal model for studying how sensory stimuli are translated into behavioral responses (Vosshall and Stocker, 2007). Indeed, highly specific survival behaviors, such as innate avoidance of stress odorants, can be traced to the activation of developmentally hardwired, single-glomerulus circuits (Stensmyr et al., 2012; Suh et al., 2004). Many of these responses are shaped by natural selection to reflect the ecological relevance of specific odorants: isoamyl acetate, a component of fermenting fruit, is typically attractive, whereas compounds such as benzaldehyde and 1-octanol are generally aversive (Knaden et al., 2012; Semmelhack and Wang, 2009). However, olfactory behavior is not entirely hardwired; the innate valence of olfactory stimuli is continuously fine-tuned throughout an animal’s life based on experience (Busto et al., 2010), specifically through dopaminergic pathways that signal reward and punishment to update memory (Liu et al., 2012; Waddell, 2010). The impact of early-life environmental exposure on adult behavior has been debated for decades (Glanzman, 2010). Early studies proposed the concept of pre-imaginal conditioning, hypothesizing that larvae reared in a scented medium could retain true associative memories into adulthood (Thorpe, 1939; Tully et al., 1994). However, subsequent behavioral reassessments demonstrated that these shifts might be driven by a persistent “chemical legacy”, residual odorants carried over from the larval environment, rather than true memory retention across metamorphosis (Barron and Corbet, 1999; Colomb et al., 2007; Ramirez et al., 2016). A central question in sensory neurobiology is how experience-dependent plasticity can persist across life stages in holometabolous insects, given that *Drosophila* undergoes radical morphological reorganization during complete metamorphosis. The adult olfactory system develops *de novo* during the pupal stage, projection neurons form protoglomeruli prior to the arrival of olfactory receptor neurons (Jefferis and Hummel, 2006; Jefferis et al., 2004), and early structural development appears largely independent of sensory input (Wong et al., 2002). Despite this developmental bottleneck and structural reorganization, sensory experiences during development and early adulthood (i.e. with an immature olfactory system) leave traces that modulate adult behavior. Extensive evidence now demonstrates that adult *Drosophila* olfactory behavior exhibits significant plasticity (Busto et al., 2010; Tully and Quinn, 1985). The antennal lobe organizes olfactory information into spatial maps where hedonic valence is represented at the output level (Hallem and Carlson, 2004; Knaden et al., 2012; Vosshall et al., 2000), and mushroom body connectivity shows structured sampling organized around natural odor relationships (Yang et al., 2023). Foundational work showed that exposing young adult flies to odorants for several days induces glomerulus-specific volumetric decreases and reduces behavioral aversion, effects attributed to central rather than peripheral adaptation (Devaud et al., 2001). Additionally, Notch signaling is required for some of the morphological and functional plasticity observed following odor exposure (Kidd et al., 2015). Crucially, this plasticity is largely restricted to a critical period during early adulthood, as older flies show little susceptibility to equivalent exposure (Devaud et al., 2003; Golovin and Broadie, 2016; Sachse et al., 2007). Beyond habituation, early odorant exposure can also act as positive reinforcement, enhancing odor-specific attraction (Arenas et al., 2009; Chakraborty et al., 2009; Twick et al., 2014), or induce latent inhibition in a mushroom body-dependent manner (Jacob et al., 2021). Furthermore, natural and diverse olfactory experiences generate greater variability and richer odor-processing patterns in the antennal lobe than homogeneous olfactory environments (Jernigan et al., 2020). At the circuit level, experience-dependent structural remodeling involves cAMP signaling and requires *rutabaga* function specifically in GABAergic local interneurons (Chodankar et al., 2020; Das et al., 2011). Recent high-resolution studies have extended these findings: prolonged exposure to ecologically relevant odors can induce dendritic expansion and volumetric increases in specific glomeruli, fundamentally altering innate aversions (Fabian and Sachse, 2023), while rearing animals with natural food odors enhances attraction through associative mechanisms, leaving early sensory representations largely stable (Dylla et al., 2023; Gugel et al., 2023). Despite these advances, the extent to which continuous developmental exposure to monomolecular odorants of varying innate valences modulates adult preference and exploratory behavior remains incompletely understood. Here, we investigated whether odorant exposure during development and early adult life, two very different developmental stages, affects olfactory-guided behavior in adult *Drosophila melanogaster*. We exposed wild-type flies to odorants of distinct innate valences by incorporating these compounds into the larval food medium. Adult flies were subsequently assessed using a two-choice behavioral assay to quantify both odor preference and general exploratory drive. Our results demonstrate that early odorant exposure significantly influences adult olfactory behavior, typically reducing aversion or enhancing attraction toward the exposed compounds, with effects that are concentration– and sex-dependent. These findings provide new insights into how non-associative developmental experiences program adult behavior in holometabolous insects.

## Materials and methods

### Strains and rearing conditions

Flies were raised in a 12h:12h light:dark (LD) cycle at 25°C in vials containing standard cornmeal medium. All experimental protocols were performed in accordance with relevant institutional ethical guidelines. Wild-type Canton-S and Berlin strains were used. To establish an unbiased baseline, stock colonies were bred on a standard medium entirely free of propionic acid for more than 10 generations prior to the experiments. For the initial validation assays (Figure 1), animals aged 3 to 7 days were used. For all subsequent choice assays, 4– to 7-day-old adult males and females were evaluated.

**Figure 1.**
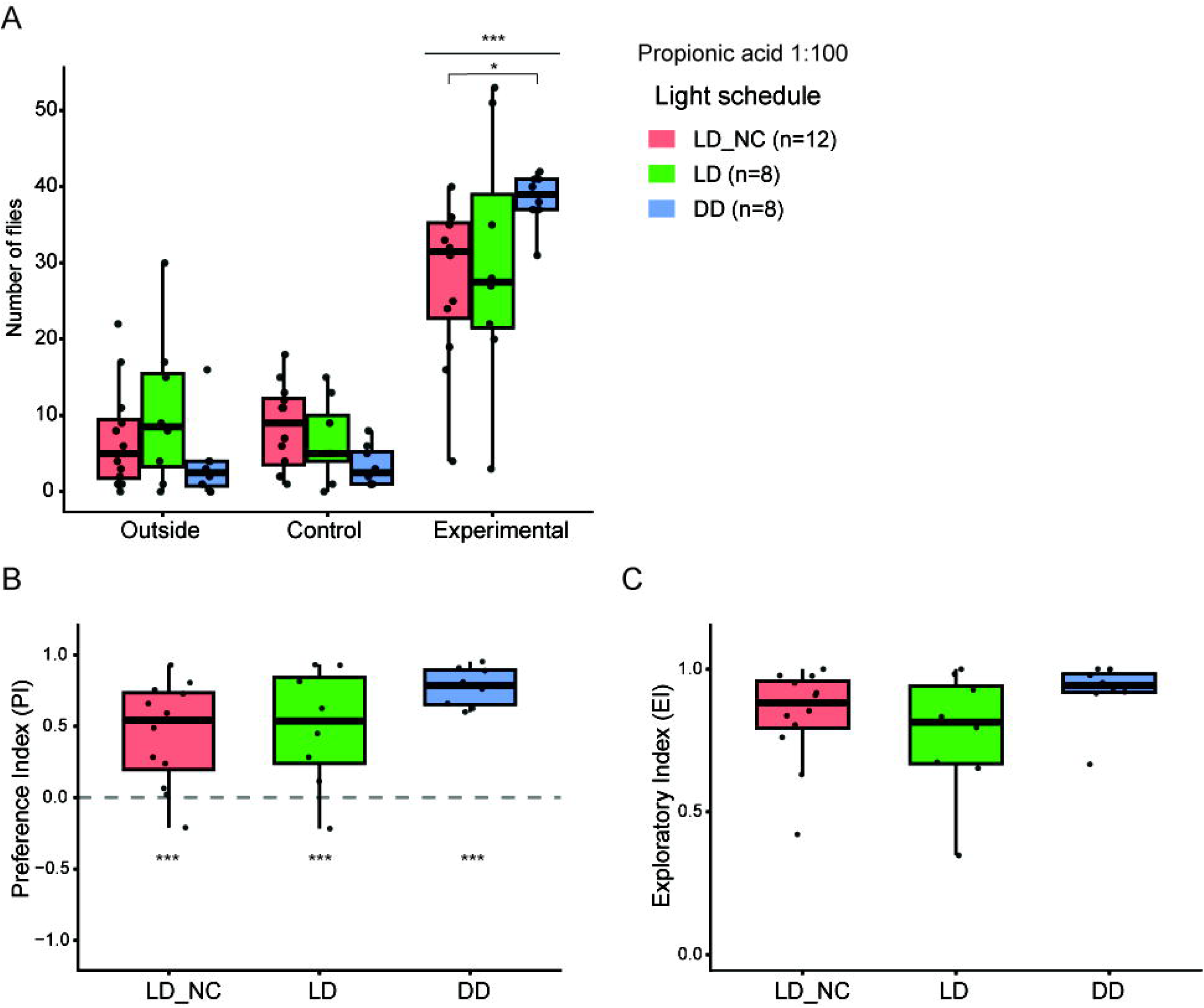
Ambient light schedules modulate the magnitude of the adult behavioral response to propionic acid. Flies were tested in a two-choice assay (propionic acid 1:100 vs. vehicle) under varying lighting conditions. **(A)** Spatial distribution. Attraction to the odorant peaked under constant darkness (DD) compared to the non-controlled light cycle (LD_NC) (interaction p < .050). **(B)** Preference Index (PI) confirms robust attraction across all lighting conditions (p < 0.001 vs. zero). **(C)** Exploratory Index (EI) indicates stable overall exploratory activity across light schedules. Boxplots display medians and interquartile ranges (IQR); dots represent individual assay arenas (n = 8-12 per condition). Asterisks indicate statistical significance from zero or between groups as determined by Type III ANOVA with Tukey’s HSD post hoc test (*** p < 0.001).

### Olfactory stimuli and developmental exposure

Monomolecular odorants were selected based on their established innate valences (Knaden et al., 2012; Semmelhack and Wang, 2009): propionic acid (appetitive), isoamyl acetate (neutral), benzaldehyde (aversive), and 1-octanol (aversive). All chemicals were obtained from Euma (Buenos Aires, Argentina). To evaluate the impact of the developmental olfactory environment, animals were reared in media supplemented with specific odorants. Odorants were incorporated directly into the liquid food substrate rather than presented passively, as active substrate interaction is critical for inducing associative learning and long-term behavioral plasticity (Dylla et al., 2023; Otárola-Jiménez et al., 2024). Initial pilot optimizations revealed that rearing larvae at the testing concentration of 1:100 resulted in high toxicity and minimal developmental survival. Consequently, to ensure robust population viability while maintaining chronic sensory exposure, the breeding medium for all the treated cohorts was supplemented at a 1:1000 dilution. It is important to note that control animals were breed in medium with no added odorant. Thus, control animals were reared on medium free of propionic acid, that is normally added to the breeding meedium, or any other odorant. For experiments where we tested the effect of propionic acid, we used a 4/1000 concentration to mimic standard *Drosophila* medium. Upon pupation, adults were cleared to precisely control emergence dates. Newly eclosed males and females (0–2 days post-eclosion) were collected and housed in sex-segregated groups of 20–30 individuals in their respective rearing media for an additional 5 days prior to testing. Control experiments indicated no behavioral differences when sexes were co-housed (data not shown).

### Behavioral two-choice assay

Adult olfactory preference was assessed using a modified trap assay (Knaden et al., 2012). Behavioral arenas consisted of custom acrylic chambers (14 × 6 × 9 cm; 756 cm³). Each chamber contained two traps made from 30 mL clear glass vials, fitted with modified yellow micropipette tips (cut to a 2 mm entry diameter). The control trap contained 200 µl of 0.1% Triton X-100 vehicle, while the experimental trap contained the test odorant diluted in the vehicle (tested at either 1:100 or 1:1000 concentrations). Between 50 and 60 flies (approximate 1:1 sex ratio) were introduced into each chamber. Assays ran for 24 hours at 25°C in complete darkness (DD), with the exception of the light-cycle validation experiments where we used different lighting conditions, a non-controlled light schedule (LD_NC), a controlled 12:12 light-dark cycle and constant darkness (Figure 1). For the LN_NC experiments were performed at laboratory bench close to a window to follow a close to natural light schedule. On the other hand, both LD and DD experiments were performed at an incubator. Multiple chambers (4 to 8) were run simultaneously in a randomized spatial arrangement, balanced equally between control and treated cohorts. Flies dead at the end of the 24-hour period were excluded from all analyses.

### Data analysis and statistics

To quantify behavior, the spatial distribution of flies (inside the experimental trap, inside the control trap, or remaining outside alive) was recorded. From these counts, two indices were calculated:

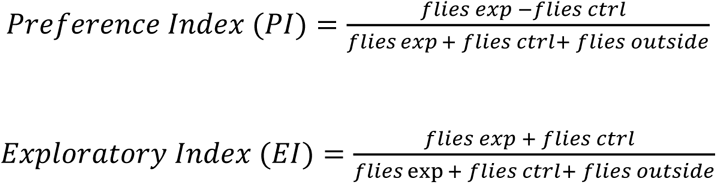

It is important to point out, that flies that stayed outside were only take into account if they were alive at the end of the experiment. The PI ranges from –1 (total avoidance) to 1 (total preference), while the EI ranges from 0 (no exploration) to 1 (maximum exploration). These metrics are established standards for evaluating *Drosophila* decision-making (Classen and Scholz, 2018; Giang et al., 2017; Knaden et al., 2012; Larsson et al., 2004; Ruebenbauer et al., 2008; Schneider et al., 2012). Statistical analyses and data visualizations were performed using R (R Core Team, 2023) within the RStudio integrated development environment (RStudio Team, 2023). Data manipulation and graphical representations were executed using the *tidyverse*, *car*, and *emmeans* packages. Animal counts were analyzed using linear models (LM) or linear mixed models (LMM, incorporating arena as a random effect) fitted for each experimental condition, modeling distribution as a function of rearing diet, trap condition, and their interaction. PI and EI values were analyzed using Type III ANOVA, evaluating the main effects of rearing diet, sex, and their interactions. Estimated marginal means were computed using ‘emmeans’, with post hoc pairwise comparisons adjusted via the Tukey HSD or Holm methods where appropriate. Deviations of PI from zero (chance level) were assessed using two-tailed one-sample t-tests. Model assumptions (normality of residuals and homoscedasticity) were rigorously verified via Shapiro-Wilk tests and visual inspection of Q-Q and residuals-vs-fitted plots. Influential observations were monitored using Cook’s distance, and no biological data points were excluded.

## Results

### Establishment of a two-choice assay to evaluate exploration and odor preference

To establish a robust two-choice olfactory behavioral assay, we first evaluated the impact of ambient lighting on odor-driven behavior using wild-type Berlin flies. Animals were reared in medium supplemented with propionic acid (4:1000) and tested against a 1:100 concentration of the same odorant. To maximize the role of the olfactory system in the behavioral task, we compared spatial distribution across three lighting schedules: a non-controlled natural light-dark cycle (LD_NC), a controlled 12:12 light-dark cycle (LD), and constant darkness (DD). A two-way ANOVA (Type III) revealed that while the main effect of lighting condition on overall distribution was not significant, F(2, 75) = 0.19, p = 0.829, a highly significant main effect of spatial choice (outside, control trap, experimental trap) was observed, F(2, 75) = 83.59, p < 0.001. Importantly, we detected a significant interaction between light schedule and trap choice, F(4, 75) = 3.13, p = 0.019, indicating that the behavioral response to propionic acid is modulated by the lighting environment (Figure 1A). Post hoc comparisons (Tukey’s HSD) showed that the number of animals entering the experimental trap was significantly higher in constant darkness compared to the LD_NC environment (p = 0.015). In contrast, lighting conditions did not affect the distribution of animals that stayed outside or entered the control trap (p > 0.05). We hypothesize that this phenomenon reflects a state-dependent shift in sensory modality weighting: in the absence of competing visual inputs, the animals’ navigational strategy becomes more heavily reliant on olfactory cues, thereby increasing the behavioral gain toward the chemical source.

To quantify these behaviors independently of absolute animal counts, we calculated a Preference Index (PI) and an Exploratory Index (EI). Analysis of the PI revealed a robust attraction to propionic acid across all experimental lighting conditions. One-sample t-tests confirmed significant positive preferences against chance level (0) for LD_NC, LD, and DD conditions (p < 0.001; Figure 1B). Subsequent comparisons indicated that the magnitude of this preference is not significantly modulated by the lighting environment (p > 0.05 for all pairwise comparisons), although a trend toward higher preference in DD compared to LD_NC was observed (p = 0.090). Finally, to evaluate if lighting conditions altered general activity levels, we analyzed the EI. This analysis confirmed that general exploratory drive remained stable across all light schedules (Figure 1C). However, similar to the spatial distribution data, the dispersion of the indices was visibly reduced under constant darkness. Taken together, these results validate the efficacy of our behavioral arena. Because constant darkness successfully maximized odor-driven trap entry and minimized behavioral variance, all subsequent experiments were conducted under DD conditions to optimally isolate olfactory navigation.

### Evaluating the role of the developmental olfactory environment on adult odor preference

To determine how the olfactory environment during development and early adulthood modulates adult behavior, we assessed responses to stimuli spanning a full spectrum of innate valences. We selected four well-characterized monomolecular odorants: propionic acid (appetitive), isoamyl acetate (neutral), benzaldehyde (aversive), and 1-octanol (aversive) (Knaden et al., 2012; Semmelhack and Wang, 2009). All subsequent choice assays utilized Canton-S wild-type flies to provide a standardized behavioral baseline for olfactory processing. Because the odorants were incorporated directly into the developmental food substrate, we hypothesized that this chronic associative exposure might act as a positive reinforcement. Consequently, utilizing innate aversive and neutral stimuli provides a robust framework to test whether continuous environmental exposure can fundamentally override or shift intrinsic hedonic valence (Dylla et al., 2023). Conversely, evaluating an innate attractant (propionic acid) allows us to determine whether an already positive valence can be further enhanced by early-life familiarity.

### Propionic acid induces moderate innate attraction and rigid exploratory behavior, lacking sexual dimorphism

To establish the baseline response to an ecologically relevant organic acid, we evaluated behavior towards propionic acid at a 1:100 testing concentration. This chemical is used as a preservative in standard *Drosophila* medium and acts as a known attractive odorant in both behavioral and calcium imaging paradigms (Das Chakraborty et al., 2022; Knaden et al., 2012). To evaluate developmental effects, animals were reared either in a medium free of propionic acid or a medium supplemented with propionic acid at a 4:1000 dilution. Testing was subsequently conducted at a higher concentration (1:100) to facilitate direct comparison with established olfactory valence maps (Knaden et al., 2012). Analysis of the spatial distribution revealed a clear preference for the experimental trap over the vehicle control, although a substantial number of animals remained outside the traps (Figure 2A). A two-way ANOVA (Type III) demonstrated a highly significant main effect of spatial choice (outside, control trap, and experimental trap) on animal distribution, F(2, 186) = 60.71, p < 0.001. However, developmental exposure to propionic acid did not alter this behavior; there was no significant main effect of the odorant presence (0 vs 4:1000) in the breeding environment, F(1, 186) = 0.05, p = 0.829, nor a significant interaction between rearing diet and spatial choice, F(2, 186) = 0.96, p = 0.380. Post hoc comparisons (Tukey’s HSD) confirmed that the number of flies entering the control trap was significantly lower than both the experimental trap and the outside zone (p < 0.001 for both), regardless of the developmental environment. Furthermore, no significant differences were observed between the number of animals entering the experimental trap and those remaining outside (p ≥ 0.480). These results indicate that the innate appetitive valence of propionic acid is highly robust and is not fundamentally increased or modulated by continuous exposure during development. To quantify the hedonic valuation of the odorant independently of spatial counts, we analyzed the Preference Index (Figure 2B). Propionic acid elicited significant attraction across all experimental groups, as confirmed by one-sample t-tests against zero (p < .001 for all groups). A two-way ANOVA revealed no significant main effects of sex (p = 0.380) or odor presence in the breeding medium (p = 0.180), and the global interaction term did not reach statistical significance, F(1, 124) = 2.31, p = 0.130. Nevertheless, planned pairwise comparisons conducted to assess specific sensitivities within sexes revealed a subtle sensitization in females: females reared on propionic acid exhibited a significant increase in preference compared to control females (p = 0.046), whereas male preference remained entirely stable regardless of their developmental environment (p = 0.890). We next evaluated general locomotor arousal by means of the Exploratory Index (Figure 2C). Strikingly, this analysis revealed a significant interaction between sex and rearing diet, F(1, 124) = 8.18, p = 0.005, indicating that developmental exposure modulates exploratory behavior differently in males and females. Post hoc analysis showed that males significantly reduced their exploratory drive when reared on propionic acid compared to control males (p = 0.008). In contrast, female exploration remained unaffected by the developmental environment (p = 0.170). Under basal control conditions, a significant sexual dimorphism was present, with males exhibiting higher exploration indices than females (p = 0.028). Crucially, the selective reduction of male exploration following developmental exposure effectively abolished this basal sexual dimorphism.

**Figure 2.**
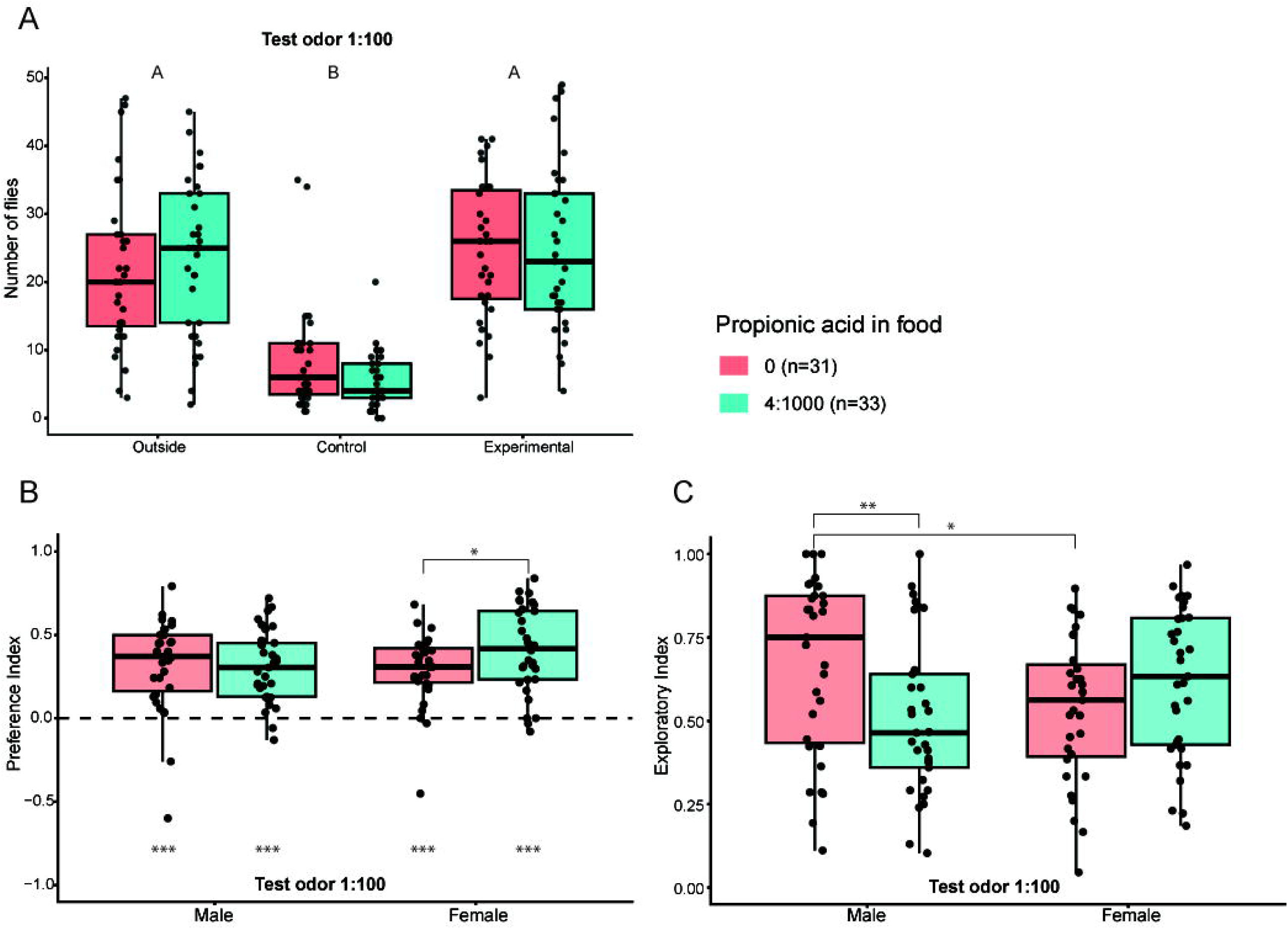
Developmental exposure to propionic acid alters adult exploration in a sex-specific manner without affecting innate preference. Flies were reared on standard medium (0) or medium supplemented with propionic acid (4:1000) and tested at a 1:100 concentration. **(A)** Spatial distribution was independent of the rearing diet (p = 0.829), maintaining robust avoidance of the control trap. **(B)** PI showed significant attraction in all groups (p < 0.001 vs. zero). Planned comparisons revealed a subtle sensitization in PA-reared females compared to control females (p = 0.046). **(C)** EI revealed a significant sex-by-diet interaction (p = 0.005). PA-reared males exhibited decreased exploration compared to control males (p = 0.008), effectively abolishing the basal sexual dimorphism observed in controls. Boxplots display medians and IQR; dots represent individual assay arenas. Letters and asterisks indicate statistically significant differences based on Type III ANOVA followed by Tukey’s HSD (** p < 0.010; *** p < 0.001; alpha = 0.05).

Taken together, our findings demonstrate that developmental exposure to propionic acid induces a sex-dependent reprogramming of adult behavior that varies according to the behavioral task. While innate olfactory preference proved remarkably robust, remaining stable in males and showing only a subtle sensitization in females, the exploratory strategy exhibited significant phenotypic plasticity. This divergence suggests that in our experimental set-up that although the neural circuits processing propionic acid valence are functionally resilient to early olfactory manipulation, sensory experience can specifically recalibrate motor reactivity and active-search thresholds in males (Figure 2). These results underscore the necessity of evaluating multiple behavioral dimensions to fully understand how early-life experiences differentially shape male and female phenotypes.

### Developmental exposure to benzaldehyde modulates mobilization and sex-specific exploration without altering odor valence

To evaluate whether developmental exposure modulates the response to an aversive stimulus, we performed the choice assay using benzaldehyde, an aversive odorant. We evaluated behavioral responses at two distinct testing concentrations (1:100 and 1:1000) to assess how developmental exposure influences decision-making under different valence contexts. As described in the methods section, the developmental breeding concentration was maintained at 1:1000 to ensure population viability. While alternative chronic delivery systems, such as constant-flow scented air, have been utilized in recent literature (Dylla et al., 2023; Fabian et al., 2023; Gugel et al., 2023), our free-choice assay allows individuals to voluntarily explore the arena before entering a trap containing the odorant or a vehicle control. In contrast to the absolute rigidity observed with propionic acid, responses to benzaldehyde revealed a high degree of behavioral plasticity. First, we analyzed the spatial distribution of animals in the choice assay. At the higher testing concentration (1:100, Figure 3Ai), benzaldehyde failed to act as a clear attractant or repellent, with no significant difference in the number of animals entering the experimental versus control trap (p > 0.300). However, a significant interaction between the developmental odorant concentration and trap distribution was observed, F(2, 87) = 5.04, p = 0.008. Flies reared on benzaldehyde (1:1000) were significantly less likely to remain outside compared to flies reared in the control medium, with no added odorant (p = 0.005), suggesting a reduction in basal aversion or neophobia. Decreasing the testing concentration to 1:1000 (Figure 3Aii) shifted the valence of benzaldehyde to attractive, eliciting a robust preference for the experimental trap (p < 0.001). Notably, the effect of the developmental odorant on induced mobilization persisted (F(2, 90) = 5.68, p = 0.005), since rearing on benzaldehyde significantly reduced the number of animals that remained outside the traps (p = 0.002), effectively maximizing overall participation in the assay. These results suggest that rearing on the odorant overcomes the basal inhibition to enter the traps, increasing overall participation, even though the entry remains non-selective between the odorant and vehicle at the higher concentration (1:100). Conversely, for the lower concentration, a significant difference in the number of animals entering the experimental trap was observed (p < 0.005 vs. Control/Outside).

**Figure 3.**
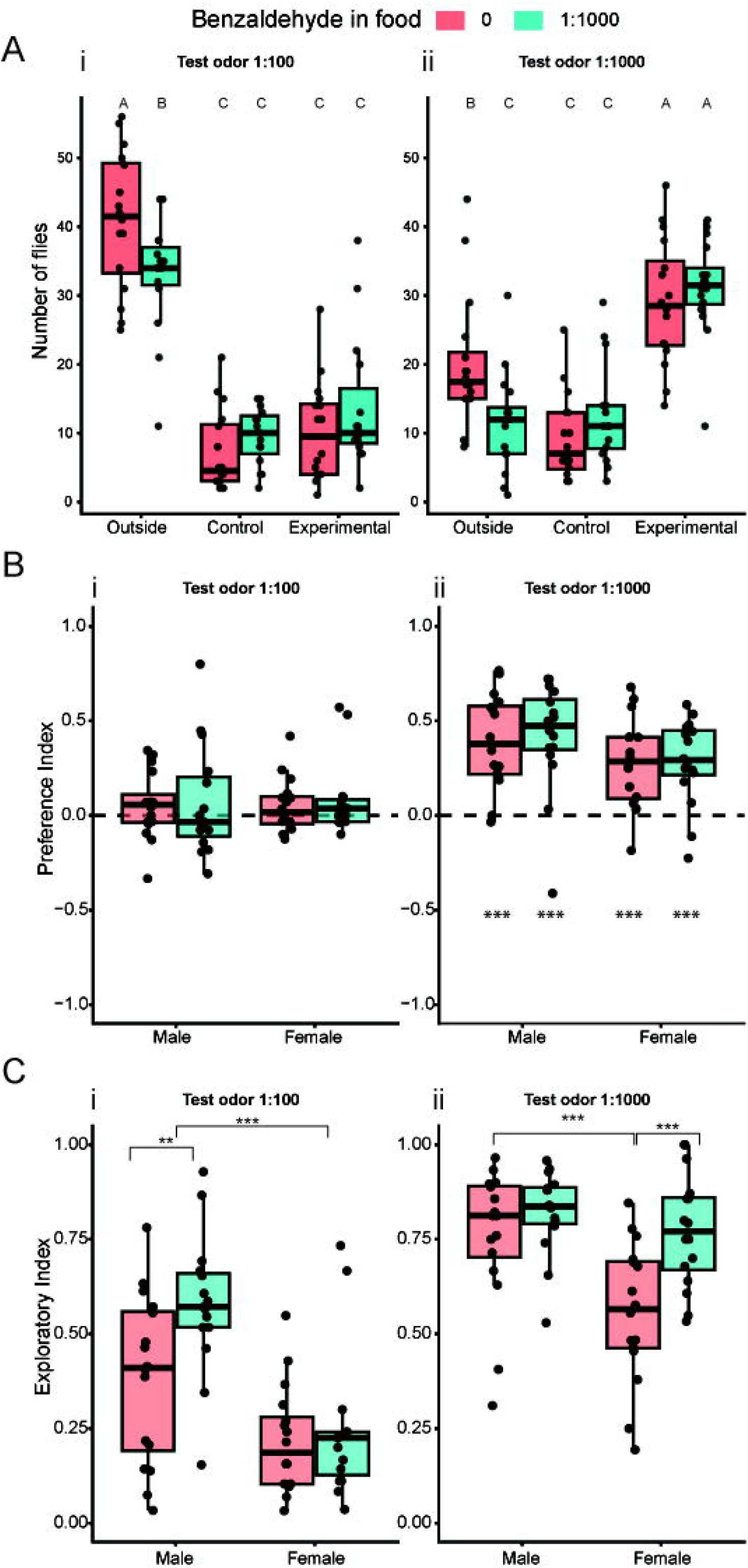
Developmental exposure to benzaldehyde modulates spatial mobilization and sex-specific exploration depending on stimulus valence. Flies were reared on standard or benzaldehyde-supplemented medium (1:1000) and tested at 1:100 or 1:1000 concentrations. **(A)** Spatial distribution. At 1:100, benzaldehyde elicited no active attraction (p > 0.300), but rearing on the odorant significantly reduced the untrapped population (p = 0.005). At 1:1000, strong attraction was observed (p < 0.001), and developmental exposure further reduced the untrapped population (p = 0.002). **(B)** PI values were neutral at 1:100 and robustly positive at 1:1000 for all groups, remaining unaltered by the developmental environment. **(C)** EI showed a concentration-dependent inversion of plasticity. At 1:100, benzaldehyde rearing specifically increased male exploration (p = 0.006). At 1:1000, rearing specifically increased female exploration (p < 0.001), abolishing the basal sexual dimorphism. Boxplots display medians and IQR; dots represent individual assay arenas. Letters and asterisks indicate statistically significant differences based on Type III ANOVA followed by Tukey’s HSD (** p < 0.010; *** p < 0.001; alpha = 0.05).

To quantify the hedonic valuation of the odorant, we analyzed the Preference Index (PI). Consistent with the distribution data, benzaldehyde at the higher testing concentration (1:100) was perceived as neutral, with PI values statistically indistinguishable from zero (p > 0.130) and unaffected by the developmental environment (p = 0.620) or sex (p = 0.920). Given that benzaldehyde has been previously characterized as an aversive stimulus (Knaden et al., 2012), the lack of distinct avoidance at this concentration was unexpected. Additionally, the lower concentration showed a clear positive preference (Figure 3Aii and Bii), becoming strongly attractive at 1:1000 (p < 0.001 vs. zero). Despite the increased trap entry observed in the spatial analysis, the magnitude of this preference was not significantly modulated by the developmental odorant exposure (p = 0.790). A marginal trend toward sexual dimorphism was noted under standard conditions, with males displaying slightly higher preference scores than females, F(1, 60) = 3.99, p = 0.050; however, this difference did not reach the threshold for strict statistical significance. These results indicate that at low concentrations and in this experimental paradigm, benzaldehyde triggers a robust innate attraction that remains resilient to developmental exposure. Early experience facilitates the decision to enter traps (mobilization) without altering the innate hedonic valence of the odorant.

Next, we examined the Exploratory Index (EI) to assess general activity levels, revealing a striking concentration-dependent inversion of sexual plasticity. At 1:100 (a neutral/mildly aversive context), rearing on benzaldehyde specifically increased exploration in males compared to controls (p = 0.006), while females remained unaffected; this effectively exacerbated the basal sexual dimorphism (males > females). In stark contrast, at the attractive concentration (1:1000), plasticity shifted to the females. While control females explored significantly less than control males (p < 0.001), rearing on benzaldehyde specifically increased female exploration (p < 0.001) to match male levels. Consequently, the sexual dimorphism observed under standard conditions was effectively abolished in the benzaldehyde-reared group (p = 0.380).

Together, these results demonstrate that developmental exposure to benzaldehyde enhances exploratory arousal in a sex-specific manner that is contingent upon the valence of the testing environment. Although the PI demonstrated that the intrinsic valence of the odorant is highly resistant to developmental exposure, early experience drastically modified mobilization dynamics. Spatial distribution analysis showed that flies reared with the odorant exhibited a strong reduction in population inertia, significantly exploring the arena to enter the traps. This odor-dependent plasticity was corroborated by the EI. Rearing on benzaldehyde profoundly increased general locomotor arousal, demonstrating that while early experience does not change the hedonic valuation of the odor, it substantially reprograms motor reactivity, revealing an olfactory plasticity mechanism that dissociates hedonic valuation from active motor search. Conversely, previous work indicates that chronic exposure to other aversive odorants, such as geosmin, reduces aversion and causes structural changes (Fabian et al., 2023). Thus, our findings emphasize that early experience reconfigures exploratory arousal and trap-entry decisions without shifting the primary sensory valuation of the stimulus, and these plasticity rules are highly odorant-dependent.

### 1-Octanol triggers concentration-dependent valence that is resilient to developmental exposure

To evaluate the stability of innate valence and its potential modulation by developmental history, we assessed the behavioral responses to 1-octanol across two distinct testing concentrations (1:100 and 1:1000). Similar to benzaldehyde, 1-octanol has been characterized as an aversive stimulus (Knaden et al., 2012); however, it elicits a relatively milder aversion compared to other canonical repellents within that established panel. In all chronic exposure groups, animals were reared on a medium supplemented with a constant 1:1000 concentration of the odorant. Analysis of the spatial distribution revealed a dramatic, concentration-dependent inversion of behavioral engagement, F(2, 114) = 71.25, p < 0.001 for 1:100; and F(2, 84) = 34.71, p < 0.001 for 1:1000. At the higher testing concentration (1:100), 1-octanol acted as a potent aversive stimulus; individuals actively avoided the experimental trap, preferentially seeking the vehicle control trap as a safe spatial alternative (p < 0.001; Figure 4Ai). Conversely, at 1:1000, the flies shifted to robust attraction, heavily populating the experimental trap over both the control trap and the “outside” zone (p < 0.001; Figure 4Aii). Crucially, these primary spatial decisions remained entirely resilient to early-life experience, as rearing history did not significantly modulate distribution patterns in either the aversive, F(2, 114) = 1.91, p = 0.150, or the attractive context, F(2, 84) = 0.94, p = 0.390.

**Figure 4.**
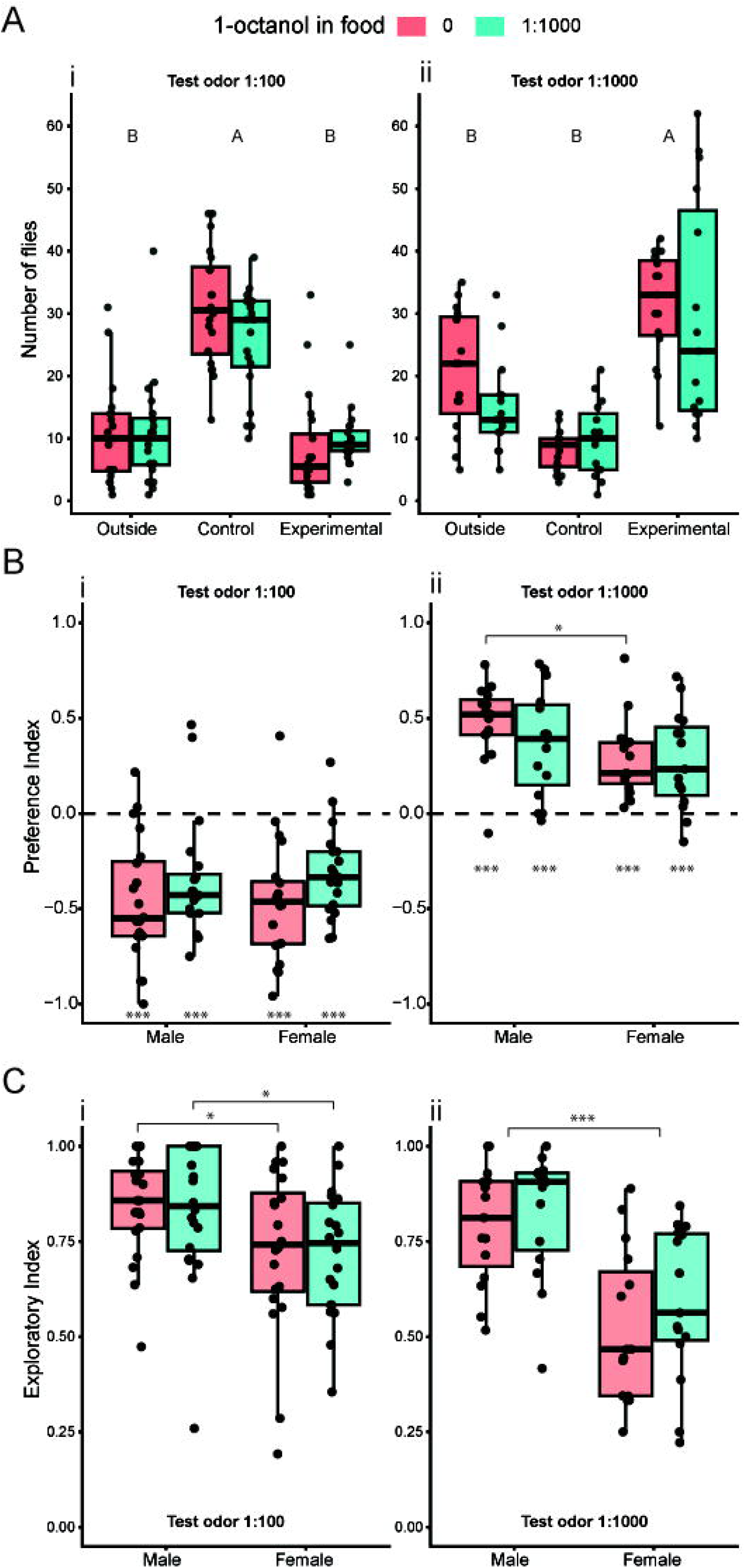
1-Octanol elicits a concentration-dependent valence reversal with resilient spatial distribution and stable exploratory drive. Flies reared on standard or 1-octanol-supplemented medium (1:1000) were tested at 1:100 and 1:1000 concentrations. **(A)** Spatial distribution. At 1:100, the odorant elicited strong active avoidance. At 1:1000, a valence reversal occurred, rendering the experimental trap highly attractive. Odorant rearing did not alter these distributions. **(B)** PI showed strongly negative values at 1:100 and strongly positive values at 1:1000 (p < 0.001 vs. zero for both). Rearing on 1-octanol induced a trend toward reduced aversion at 1:100 (p = 0.066) and abolished the basal sexual dimorphism at 1:1000 (p = 0.320). **(C)** EI remained unaffected by developmental exposure across all contexts (p > 0.290). A strong basal sexual dimorphism (males > females) persisted regardless of rearing diet. Boxplots display medians and IQR; dots represent individual assay arenas. Letters and asterisks indicate statistically significant differences based on Type III ANOVA followed by Tukey’s HSD (*** p < 0.001; alpha = 0.05).

To further quantify these shifts in valence, we analyzed the Preference Index (PI). At 1:100, all groups exhibited strongly negative PI values (p < 0.001 vs. zero), confirming the innate aversion observed in the distribution data (Figure 4Bi). However, rearing on 1-octanol induced a biologically relevant trend toward habituation; treated flies displayed a globally less negative PI compared to controls, suggesting a subtle desensitization (p = 0.066). At 1:1000, all groups showed strongly positive PI values (p < 0.001 vs. zero). Interestingly, the baseline sexual dimorphism observed in control-reared animals, where males display higher attraction than females (p = 0.033), was sensitive to developmental exposure. Rearing on 1-octanol (1:1000) induced a male-specific trend toward reduced attraction (p = 0.200) which was sufficient to completely abolish the sexual dimorphism in the treated cohort (p = 0.320).

Finally, we examined the Exploratory Index (EI) to determine if general behavioral drive was influenced by developmental exposure. Unlike the subtle plasticity observed in odor preference, exploratory behavior was entirely rigid against early odorant exposure. A strong, basal sexual dimorphism (males > females) persisted across both the 1:100 (p < 0.050) and 1:1000 (p < 0.001) contexts (Figure 4C). Rearing on 1-octanol (1:1000) had no significant effect on overall exploration levels across any group (p > 0.290). These results indicate that while early experience can finely tune sensory valuation through habituation, the broad exploratory drive and the primary spatial navigation decisions for 1-octanol are governed by inflexible, innate motor programs.

### Isoamyl acetate triggers a fixed behavioral program characterized by innate attraction and high motor inertia

To determine whether the plasticity rules observed with the previous odorants generalize to other olfactory cues, we assessed the behavioral responses to isoamyl acetate. Although this odorant has been previously categorized as neutral (Knaden et al., 2012), its reported valence score places it at the extreme lower boundary of this category, positioning it functionally close to mild repellents like 1-octanol. As in previous experiments, animals were reared continuously on a medium supplemented with a 1:1000 concentration of the odorant.

Spatial distribution analysis revealed a conflict between innate attraction and high baseline inertia across both testing concentrations. At 1:100, while flies significantly preferred the experimental trap over the control (p = 0.012), a substantial portion of the population remained outside. The number of animals remaining outside was statistically indistinguishable from those entering the experimental trap (p = 0.410; Figure 5Ai). This behavioral inertia was even more pronounced at the 1:1000 concentration, where the fraction of the population remaining outside was significantly larger than the fraction entering the experimental trap (p < 0.050). Crucially, developmental exposure to isoamyl acetate did not modulate this spatial distribution; two-way ANOVAs confirmed the absolute rigidity of this behavior, revealing no significant main effect of rearing diet at either 1:100 (p = 0.480) or 1:1000 (p = 0.400).

**Figure 5.**
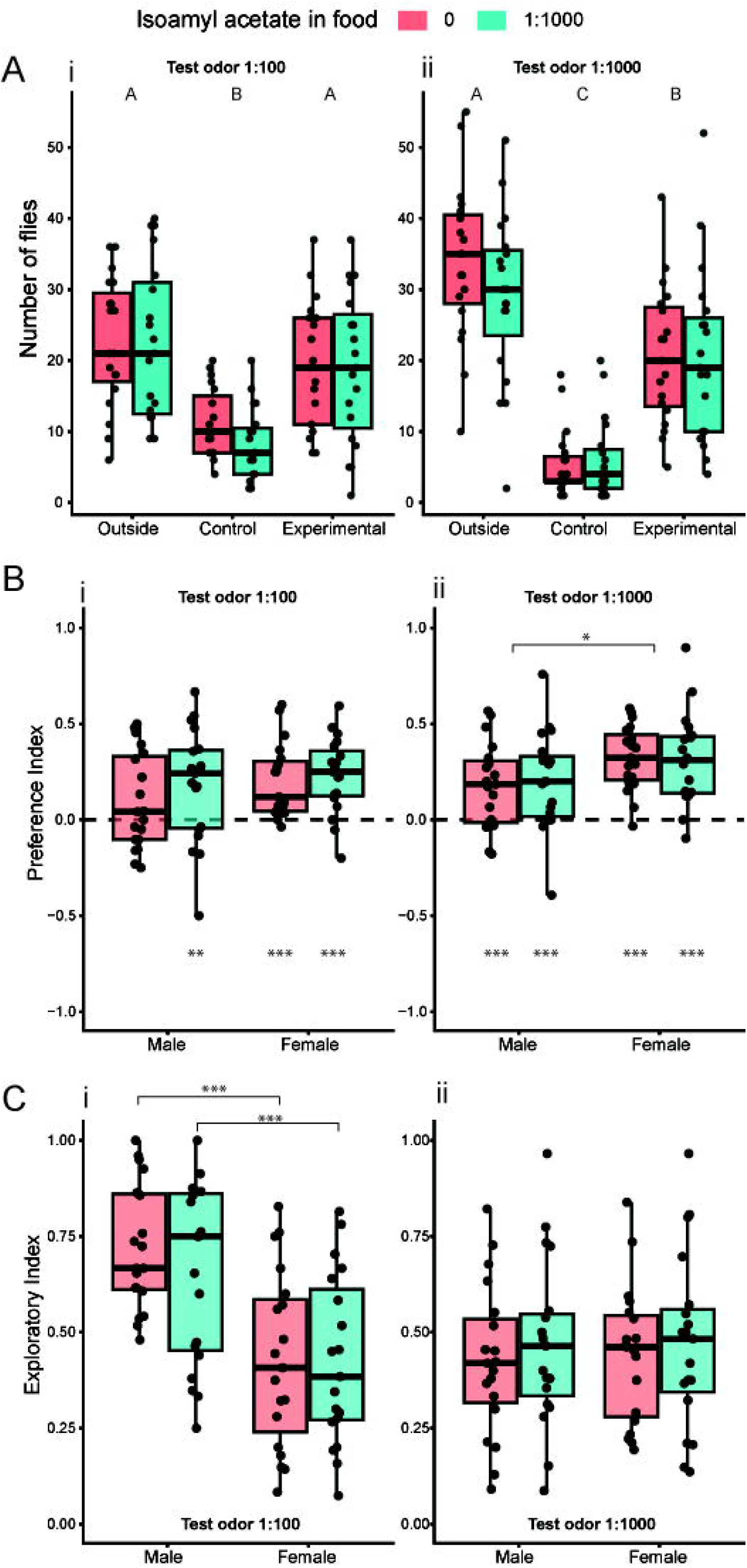
Isoamyl acetate triggers an innate, concentration-dependent attraction characterized by high baseline inertia and rigid exploratory drives. Flies reared on standard or isoamyl acetate-supplemented medium (1:1000) were tested at 1:100 and 1:1000 concentrations. **(A)** Spatial distribution. The odorant acted as an innate attractant at both concentrations (Experimental > Control). However, high baseline inertia was observed, with the outside zone retaining a substantial population. This distribution remained unaffected by developmental exposure. **(B)** PI exhibited positive values at 1:100 (p < 0.010) and 1:1000 (p < 0.001). Developmental exposure left odor preference completely intact. **(C)** EI was unaffected by developmental exposure at both concentrations (p > 0.470). The basal sexual dimorphism (males > females) present at 1:100 was absent at 1:1000, independent of the rearing diet. Boxplots display medians and IQR; dots represent individual assay arenas. Letters and asterisks indicate statistically significant differences based on Type III ANOVA followed by Tukey’s HSD (** p < 0.010; *** p < 0.001; alpha = 0.05).

The stability of this innate response was further corroborated by quantifying hedonic valence through the Preference Index (PI). At 1:100, all groups exhibited a significant but mild attraction against zero (p < 0.010), with the exception of control males, which showed only a marginal trend toward attraction. Decreasing the testing concentration to 1:1000 elicited a stronger, highly significant attraction across all groups (p < 0.001) and revealed a basal sexual dimorphism wherein females were more attracted than males (main effect of sex, p = 0.014; Figure 5Bii). However, in stark contrast to the habituation trends observed with 1-octanol and the robust mobilization plasticity induced by benzaldehyde, rearing on isoamyl acetate induced no plastic changes in odor preference at either testing concentration (p = 0.280 for 1:100; p = 0.920 for 1:1000).

Finally, analysis of the Exploratory Index (EI) confirmed that general motor arousal in response to isoamyl acetate is strictly hardwired. Exploratory activity exhibited a strong basal sexual dimorphism at the 1:100 concentration, with males exploring significantly more than females (main effect of sex, p < 0.001). Interestingly, this dimorphism was completely abolished at the lower 1:1000 concentration (main effect of sex, p = 0.810; pairwise contrasts, p > 0.800), driven primarily by a reduction in male exploration. Crucially, regardless of the testing concentration or the presence of baseline sexual dimorphisms, developmental exposure to the odorant failed to induce any plastic changes in general exploratory behavior across all tested groups (main effect of rearing diet, p = 0.490 for 1:100; p = 0.470 for 1:1000). Thus, the behavioral programs governing both hedonic preference and general exploration for this odorant are entirely resistant to early-life sensory manipulation.

### Global summary of phenotypic plasticity

Taken together, these results demonstrate that behavioral plasticity induced by early olfactory experience is not a global or generalized phenomenon, but rather a finely tuned process dependent on the identity, valence, and concentration of the odorant. Our stimulus panel revealed a full spectrum of sensorimotor adaptations. On one end, isoamyl acetate triggers a rigidly innate behavioral program, where both hedonic preference and exploratory drive remain entirely impervious to manipulation of the olfactory environment. Propionic acid similarly maintains a rigid innate attraction, but selectively recalibrates male exploratory drive. On the other end, odorants like benzaldehyde and 1-octanol expose highly specific and divergent plasticity mechanisms. While developmental exposure to benzaldehyde reveals a functional dissociation, strongly reprogramming spatial mobilization and motor arousal without altering the intrinsic valence of the odor, 1-octanol illustrates a fine-tuning of the valence itself, inducing trends of sensory habituation capable of suppressing basal sexual dimorphisms without modifying general locomotor programs. In conclusion, the olfactory system employs tailor-made adaptive strategies in response to chronic early-life exposure, ranging from absolute behavioral inflexibility to the selective modulation of exploratory drive or hedonic processing, highlighting a remarkable complexity in developmental sensorimotor integration.

## Discussion

Our study provides a comprehensive characterization of how chronic olfactory exposure during critical developmental windows and early adulthood shapes adult behavior in the fruit fly *Drosophila melanogaster*. We demonstrate a fundamental functional dissociation: while the core neural circuits governing innate hedonic valence remain remarkably resilient to environmental manipulation, early-life experience profoundly reprograms exploratory arousal and motor search strategies. This modularity in behavioral plasticity suggests that *Drosophila* prioritizes the stability of essential sensory-driven signals while maintaining the flexibility to adapt its navigational intensity to a persistent olfactory landscape.

### Methodological optimization and sensory salience

The success of our behavioral paradigm relied on the rigorous calibration of environmental parameters to isolate olfactory-driven behavior from competing sensory modalities. We found that constant darkness (DD) provided the most robust and least variable response to propionic acid compared to non-controlled light cycles (LD_NC). This peak in attraction under DD likely reflects a state-dependent shift in sensory modality weighting. In the absence of visual inputs, the animal’s navigational strategy becomes more heavily reliant on chemical cues, thereby increasing the behavioral gain toward the odor source. Furthermore, we identified a critical physiological bottleneck regarding CO2 anesthesia. Brief exposure followed by a mandatory 24-hour recovery period was essential to prevent confounding motor deficits and high mortality. Animals tested before complete recovery exhibited significantly lower appetitive indices, suggesting that the metabolic cost of recovery may interfere with sensory perception or the motricity required to enter the trap vials. This underscores the necessity of accounting for metabolic state when evaluating population-level dynamics in choice assays.

### Resilience of innate valence: an ecological strategy

Contrary to our initial hypothesis, incorporating odorants directly into the larval feeding substrate did not induce a universal appetitive shift. While pre-imaginal conditioning has been observed in other insects, such as *Apis mellifera* (Arenas et al., 2009; Ramirez et al., 2016), *Drosophila* appears to possess a more rigid olfactory architecture. This resilience of innate valence likely reflects the species’ biology as a generalist. *Drosophila melanogaster* exploits a wide variety of resources; thus, maintaining robust innate preferences prevents maladaptive specialization to a single niche, even when the developmental environment is saturated with a specific compound. While previous studies indicate that third-instar larvae can form aversive memories (Fabian et al., 2023), our appetitive developmental exposure lacked the same transformative power over adult preferences. This suggests that the circuits governing innate attraction, particularly for ecologically relevant organic acids like propionic acid, are more deeply hardwired than those mediating avoidance (Knaden et al., 2012; Vosshall and Stocker, 2007). This resilience aligns with recent findings that under naturalistic odor concentrations, the olfactory system prioritizes functional stability over structural remodeling, ensuring that critical ecological cues remain reliably encoded (Gugel et al., 2023).We hypothesized that evolutionarily, the cost of misidentifying a toxic resource is significantly higher than the benefit of optimizing an already attractive stimulus, leading to a “safety-first” encoding of valence.

### Sex-specific exploratory plasticity and circuit implications

One of the most striking findings was the selective reconfiguration of exploratory dynamics. In propionic acid-reared animals, we observed a significant sex-by-diet interaction, where males selectively reduced their exploratory activity matching female levels, effectively abolishing the basal sexual dimorphism observed in control cohorts. Similarly, developmental exposure to benzaldehyde did not shift its intrinsic valence but drastically modified mobilization. Flies reared with the odorant showed a significant reduction in population inertia, with more individuals voluntarily entering the traps regardless of the concentration tested. This dissociation suggests that while early sensory experience leaves primary odor representations in the antennal lobe largely stable (Gugel et al., 2023), it specifically recalibrates search thresholds in higher-order integration centers. At the circuit level, repeated odor exposure has been shown to modulate ongoing spontaneous activity in the antennal lobe, significantly reducing background noise to facilitate more robust and highly specific odor representations (Franco and Yaksi, 2021). This enhanced signal-to-noise ratio could explain the reduction in population inertia, allowing flies to optimize their motor search strategies toward familiar stimuli without altering the underlying hedonic value. Furthermore, plasticity may occur at the level of the lateral horn, which supports innate valence coding (Das Chakraborty et al., 2022), or within the mushroom body-dependent circuits that regulate motor reactivity (Jacob et al., 2021). Indeed, specific compartments within the mushroom body have been shown to encode discrete representations of novelty versus familiarity (Hattori et al., 2017). Olfactory experience is also known to skew the synaptic balance of mushroom body output neurons (MBONs) to steer behavioral choice and drive the individualization of olfactory coding (Owald and Waddell, 2015). Such plasticity at the MBON level could serve as the primary mechanism driving the familiarization-dependent shifts in motor arousal observed in our assays.

### Physiological constraints and future directions

A primary limitation was the method of chronic exposure. Incorporating high concentrations of odorants directly into the food often resulted in excessive toxicity and reduced larval survival. Indeed, exposing the olfactory system to intense, unnatural odor concentrations might force a loss of receptor binding specificity that does not reflect the fly’s natural ecology. Furthermore, recent evidence demonstrates that under naturalistic, low-concentration conditions, the olfactory circuit actively prioritizes functional stability over massive structural remodeling to ensure that critical ecological cues remain reliably encoded (Gugel et al., 2023). Future work should explore alternative delivery systems, such as constant-flow scented air (Dylla et al., 2023), to reach higher exposure thresholds without inducing physiological stress. Another technical possibility could be to use and intermittent stimulation regime to avoid potential adaptation. Additionally, investigating learning-deficient mutants like *rutabaga* and *dunce* will be essential to determine if this reprogramming of exploratory arousal depends on canonical cAMP-signaling pathways involved in neuronal plasticity (Chodankar et al., 2020).

### The ecological context of memory consolidation

While our data demonstrates a decoupling of exploratory arousal from hedonic valence, the persistence of these behavioral shifts in a natural environment likely depends on the specific ecological context of the exposure. Recent behavioral paradigms reveal that female *Drosophila melanogaster* will significantly increase their preference for a previously experienced substrate, but this requires that the flies not only smell or touch the substrate, but actually oviposit on it (Otárola-Jiménez et al., 2024). The physical act of egg-laying serves as a crucial reinforcing stimulus, resulting in a robust long-term memory that alters oviposition preference for at least four days (Otárola-Jiménez et al., 2024). Indeed, odor-regulated oviposition behavior is a major driver of ecological specialization (Álvarez-Ocaña et al., 2023), and learning-based oviposition constancy allows insects to efficiently exploit familiar resources (Nataraj et al., 2021). Furthermore, environmental stressors encountered during this period, such as the detection of predators, can induce non associative long-term memories that drastically alter subsequent egg-laying decisions (Kacsoh et al., 2015a; Kacsoh et al., 2015b). Therefore, the extent to which early olfactory experience stably reconfigures adult choice is likely gated by whether the exposure is reinforced by critical life-history events like feeding or oviposition.

### Genetic background and strain-specific variability

Finally, it is essential to contextualize these sensory adaptations within the genetic framework of the tested population. While our study used the classical Canton-S strain to provide a standardized behavioral baseline, host odor preference and selectivity are known to be highly dependent on the genotypic background. Systematic comparisons across different *D. melanogaster* wild-type lines (such as Canton-S versus wild-type Berlin) have demonstrated significant and repeatable variations in their behavioral responses, thresholds, and overall responsiveness to optimal and suboptimal olfactory stimuli (Ruebenbauer et al., 2008). This is further supported by evidence that the underlying genetic architecture of olfactory behavior varies significantly among different wild-derived populations (Lavagnino et al., 2008). Moreover, recent comparative studies demonstrate that the valence of specific odorants is not universally fixed across the *Drosophila* genus, but is instead shaped by evolutionary adaptations at multiple processing levels to match the unique ecological niche of each species (Depetris-Chauvin et al., 2023). Consequently, the observed dissociation between motor arousal and hedonic valence may exhibit phenotypic variability across different wild populations, reflecting the diverse evolutionary strategies flies use to navigate dynamic, unpredictable olfactory environments. Furthermore, even within isogenic populations, stochastic developmental variations and micro-environmental differences are known to generate profound behavioral individuality (Mollá-Albaladejo and Sánchez-Alcañiz, 2021). Thus, early-life olfactory experience likely interacts with both evolutionary hardwiring and individual developmental trajectories to optimize motor search strategies.

## Acknowledgements

The authors wish to thank all the members of the Sensory Physiology and Plasticity laboratory for fruitful discussions of this work and Esteban Beckwith for comments on this manuscript. F.F.L. and N.P. are members of the Argentina National Research Council (CONICET). L.A.D. is funded by a CONICET doctoral fellowship.

## Author contributions

*Conceptualization:* F.F.L., N.P.; *Methodology:* M.A., F.F.L., N.P.; *Formal analysis:* M.A., L.A.D., N.P.; *Investigation:* M.A., L.A.D, N.P.; *Resources:* F.F.L., N.P.; *Data Curation:* M.A., N.P.; *Writing – original draft:* N.P.; *Writing – review and editing:* M.A., L.A.D., F.F.L., N.P.; *Visualization:* L.A.D, N.P.; *Supervision:* N.P. *Funding acquisition:* F.F.L, N.P.

## Data availability

The data set supporting the results of this article are included within the article.

